# Memory control deficits in the sleep-deprived human brain

**DOI:** 10.1101/2023.11.07.565941

**Authors:** Marcus O. Harrington, Theodoros Karapanagiotidis, Lauryn Phillips, Jonathan Smallwood, Michael C. Anderson, Scott A. Cairney

## Abstract

Sleep disturbances are associated with intrusive memories, but the neurocognitive mechanisms underpinning this relationship are poorly understood. Here, we show that an absence of sleep disrupts prefrontal inhibition of memory retrieval, and that the overnight restoration of this inhibitory mechanism is predicted by time spent in rapid eye movement (REM) sleep. The functional impairments arising from sleep loss are linked to a behavioural deficit in the ability to suppress unwanted memories, and coincide with a deterioration of deliberate patterns of self-generated thought. We conclude that sleep deprivation gives rise to intrusive memories via the disruption of neural circuits governing mnemonic inhibitory control, which may rely on REM sleep.

## Introduction

Memories of unpleasant experiences can intrude into conscious awareness, often in response to reminders. While such intrusive memories are an occasional and momentary disturbance for most people, they can be recurrent, vivid and upsetting for individuals suffering from mental health disorders such as depression, anxiety and post-traumatic stress disorder^1^. Given the transdiagnostic significance of intrusive memories, a better understanding of the mechanisms that precipitate their occurrence is vital to improving emotional wellbeing and reducing the global burden of mental illness.

One way that people ward off intrusive memories is by suppressing their retrieval, purging them from awareness. Direct suppression of unwanted memories serves an adaptive function in that it weakens the corresponding memory trace and thereby decreases the likelihood of future memory intrusions^2–8; for a review, see: 9^. In previous work, we showed that adaptive memory suppression is critically dependent on sleep^10^. Whereas well-rested individuals were able to override retrieval operations and reduce the emergence of intrusive memory content in subsequent trials, sleep-deprived individuals failed to suppress the target memories effectively, which remained intrusive over time. Moreover, the adaptive benefits of memory suppression were predicted by high-frequency heart rate variability (HF-HRV; 0.15-0.40 Hz)—a physiological correlate of cognitive control^11,12^—in sleep-rested but not sleep-deprived individuals, suggesting that mnemonic control processes linked to HF-HRV may be disrupted by an absence of sleep.

At the neurocognitive level, memory suppression is orchestrated by right dorsolateral prefrontal cortex (rDLPFC), which, via top-down inhibitory modulation, downregulates retrieval operations in hippocampus^2,3,13,14^, a process that relies on local GABAergic inhibiton^15^. We recently proposed that disruption to this memory control network by sleep deprivation can explain how recurrent failures of memory suppression arise after prolonged wakefulness^16^. Indeed, rDLPFC is a domain-general inhibitory control region implicated in the stopping of actions as well as unwanted memories^13,17^; sleep deprivation reduces rDLPFC engagement during motor response inhibition, leading to a breakdown in task performance^18,19^. Relatedly, sleep deprivation impairs prefrontal control of amygdala during threat-related information processing, prompting an overnight increase in state anxiety^20^. Whether an absence of sleep impairs prefrontal inhibition of hippocampus during memory suppression has yet to be determined, but is key to understanding the neurocognitive mechanisms by which sleep problems give rise to intrusive thoughts.

Suppressing unwanted memories in a goal-directed manner is also broadly dependent on the adaptive segregation of whole-brain functional networks. The default mode network (DMN) is a collection of interacting brain regions, including medial prefrontal and posterior cingulate cortex, that are commonly activated at rest (i.e., when focusing on one’s internal mental state) and deactivated during attention-demanding tasks^21,22^. The DMN is anti-correlated with the frontoparietal cognitive control network (CCN), a task-positive network encompassing lateral prefrontal and superior parietal areas^23^. Supported by ascending arousal input from the thalamus, performance of externally driven, goal-directed tasks is thought to rely on disengagement of the DMN and reciprocal engagement of the CCN^24,25^. Inappropriate perseverance of the DMN during attention-demanding cognitive tasks, as well as unstable thalamic activity, are a common consequence of sleep deprivation^26,27^. Likewise, resting-state fMRI data derived from sleep-deprived individuals reveals a breakdown of functional decoupling between the DMN and CCN^23,28^ and aberrant connectivity between the DMN and thalamus^29^, potentially giving rise to failures of adaptive memory control.

Disruption to the brain networks underlying dynamic cognitive control by sleep deprivation should manifest behaviourally not only in failures of adaptive memory suppression, but also in the tendency to engage in fewer deliberate, on-task thoughts. Indeed, when assessed in externally demanding task contexts, sleep-deprived individuals report a higher proportion of task-unrelated thoughts than well-rested people^30^, signifying a breakdown of control processes that allocate attentional resources in accordance with environmental requirements. However, extant findings concerning the effects of sleep loss on self-generated mental content are based on the self-reported categorisation of on-task and off-tasks thoughts. Whether phenomenological patterns of ongoing thought following sleep deprivation are consistent with a degradation of cognitive control has yet to be established.

We sought to delineate the neurocognitive mechanisms through which sleep deprivation gives rise to intrusive memories and the associated consequences for ongoing patterns of self-generated thought. First, we tested the hypothesis that sleep deprivation impairs prefrontal inhibition of memory retrieval operations in hippocampus. Participants engaged in memory suppression while undergoing functional magnetic resonance imaging (fMRI) after one night of total sleep deprivation or restful sleep. We tested the prediction that sleep deprivation weakens rDLPFC engagement during memory suppression, resulting in a concomitant increase in hippocampal activation and a failure to adaptively downregulate intrusive memories.

Given the deleterious effects of sleep deprivation on memory suppression, a reciprocal question concerns the specific components of sleep that underpin the overnight restoration of prefrontal mnemonic control. Rapid eye movement (REM) sleep is thought to play an important role in decreasing next-day brain reactivity to emotional experiences^31^, and might similarly support the engagement of top-down control processes when confronted with reminders to unwanted memories. Indeed, individuals suffering from psychiatric mood disorders who have difficulty suppressing unsolicited thoughts often exhibit abnormalities of REM sleep^32–36^. We therefore recorded restful sleep with polysomnography to determine the contributions of REM sleep to memory suppression processes governed by rDLPFC.

We also sought to confirm prior findings that sleep deprivation impairs the adaptive segregation of brain networks governing internally focused mental processing and externally directed cognitive control. Sleep-rested and sleep-deprived participants thus completed a resting-state fMRI scan immediately after the memory suppression task, allowing us to test the prediction that sleep deprivation impairs functional decoupling between the DMN and CCN, and reduces thalamic connectivity with these networks.

Finally, we asked whether sleep deprivation reduces the occurrence of spontaneous thoughts corresponding to task-related mental content. Before and after the overnight delay, participants described the content of their ongoing thoughts as they engaged in either a cognitively demanding task that required engagement of working memory (1-back) or a non-demanding task (0-back). This allowed us to test the prediction that sleep deprivation reduces patterns of deliberate, on-task thinking, consistent with a breakdown of cognitive control.

Supporting our hypothesis, sleep deprivation markedly reduced rDLPFC engagement during memory suppression, leading to failures of adaptive memory control. Among sleep-rested individuals, the magnitude of suppression-related activity in rDLPFC was correlated with time spent in REM sleep, consistent with a role for REM sleep in the overnight restoration of prefrontal memory suppression mechanisms. In keeping with prior work, sleep deprivation increased resting-state functional connectivity between the DMN and CCN, but reduced connectivity between the DMN and thalamus. Finally, sleep loss led to a deterioration of deliberate, on-task thinking, irrespective of external cognitive demands. These findings support a framework in which sleep deprivation gives rise to intrusive and uncontrolled thoughts via a breakdown of the neural circuitry underpinning mnemonic inhibitory control^16^.

## Results

### Sleep deprivation impairs prefrontal memory control

After a night of sleep deprivation (n=43; 18 male; mean ± SD age: 19.58 ± 1.72 years) or restful sleep (n=42; 12 male; mean ± SD age: 20.33 ± 2.43 years), participants entered an MRI scanner and performed the Think/No-Think (TNT) task wherein they attended to reminder cues (faces) that were each paired with an emotionally negative or neutral scene. For each reminder cue, participants either actively retrieved the corresponding scene (Think trials) or directly suppressed its retrieval (No-Think trials), and then indicated whether the reminder had evoked awareness of the scene (allowing us to isolate failed suppression attempts, referred to hereafter as intrusions). Experimental procedures are illustrated in Fig. 1.

**Figure 1.**
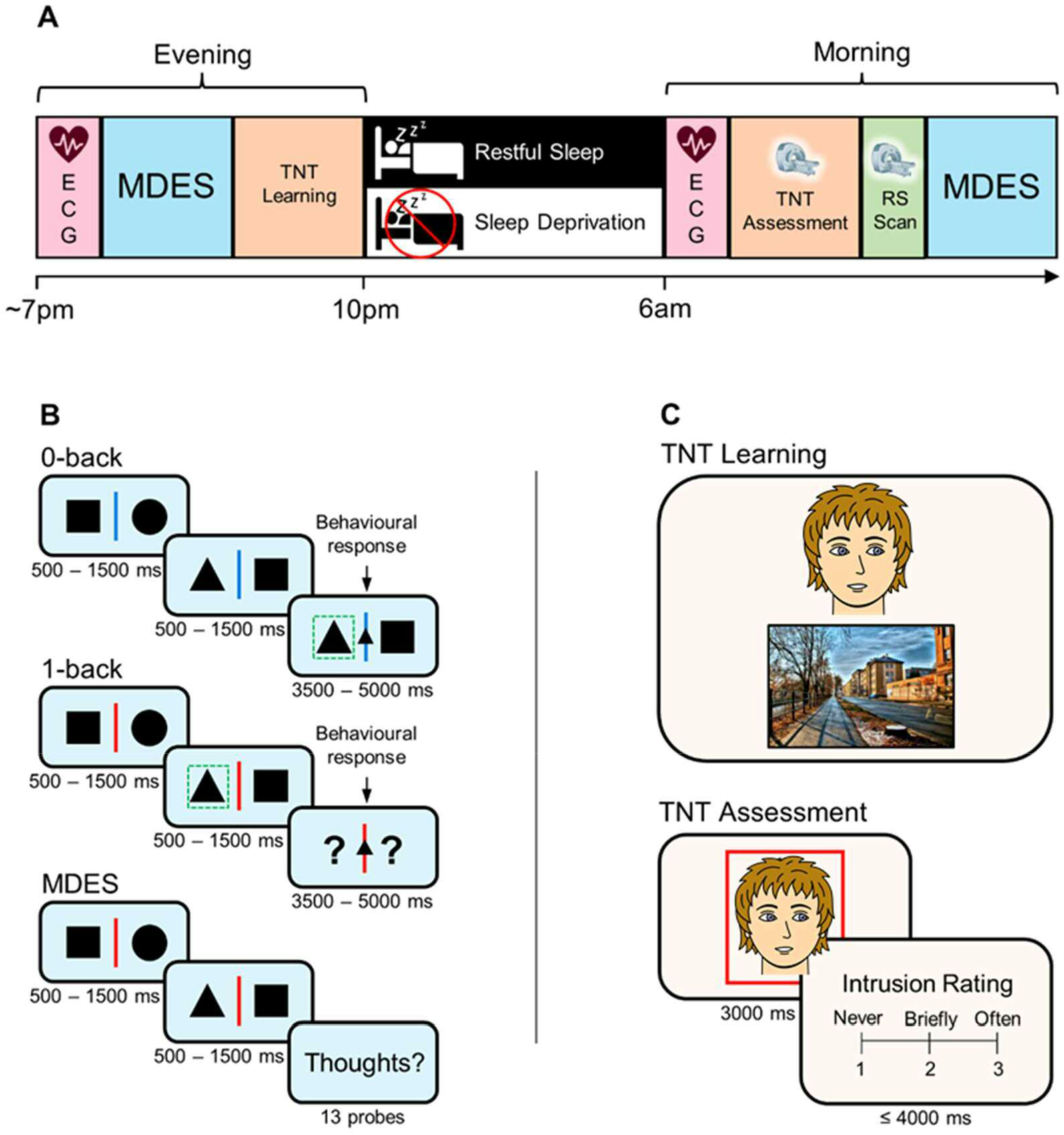
Experimental procedure. (A) Timeline. The evening session began with a resting electrocardiography (ECG) recording. Participants then completed the first of two multidimensional experience sampling (MDES) tasks and the Think/No-Think (TNT) learning phase, before either sleeping overnight in the laboratory where their sleep was recorded with polysomnography (PSG; restful sleep group) or remaining awake throughout the night (sleep deprivation group). The morning session began with another resting ECG recording. Participants then completed the TNT assessment phase inside a magnetic resonance imaging (MRI) scanner, after which a resting-state (RS) scan was acquired. Finally, participants completed the MDES task again. (B) MDES task. Participants monitored pairs of shapes. After 2-5 of these non-target trials, a target trial occurred, wherein an additional shape appeared in the centre of the screen, prompting participants to press a button corresponding to the side of the screen that the matching shape appeared on the present trial (0-back) or the immediately preceding trial (1-back). Occasionally, instead of a target trial, participants were required to indicate via a rating scale (1-10) the extent to which the contents of their ongoing thoughts matched a series of 13 thought probes. (C) TNT task. In the TNT learning phase, participants memorised 48 face-scene pairs. In the TNT assessment phase, faces were shown in isolation inside red or green frames. For red-framed faces (No-Think trials), participants were instructed to suppress (i.e., avoid thinking about) the associated scene. For green-framed faces (Think trials; not shown), participants were instructed to visualize the associated scene. After each trial, participants reported the extent to which they thought about the paired scene (never, briefly or often).

#### Behaviour

We first assessed the impact of sleep deprivation on behavioural expressions of memory control. Repeatedly suppressing intrusive memories renders them less intrusive over time^2–5,7,8^. Replicating this finding, participants showed overall decreasing numbers of intrusions (i.e., instances in which No-Think trials triggered awareness of the associated scene) across trial blocks (*F*(3.04, 218.78)=19.68, *p*<.001, *η_p_^2^*=0.21). This adaptive benefit of suppression was affected by group membership (*F*(3.04, 218.78)=2.92, *p*=.035, *η_p_^2^*=0.04; Table 1), suggesting that sleep deprivation altered the rate at which suppressed memories became less intrusive over time. Indeed, sleep-deprived participants were significantly impaired in their ability to suppress unwanted memories over TNT trial blocks relative to sleep-rested participants, as indicated by lower intrusion slope scores (see Methods; *W*=872, *p*=.043, *r_rb_*=0.27; Fig. 2A). Adaptive suppression was unaffected by the emotional valence of the target scenes (negative or neutral; *F*(4, 288)=0.60, *p*=.66, *η_p_^2^*<0.01), and scene valence had no impact on the adaptive suppression deficit among sleep-deprived participants (*F*(4, 288)=1.34, *p*=.26, *η_p_^2^*=0.02).

**Figure 2.**
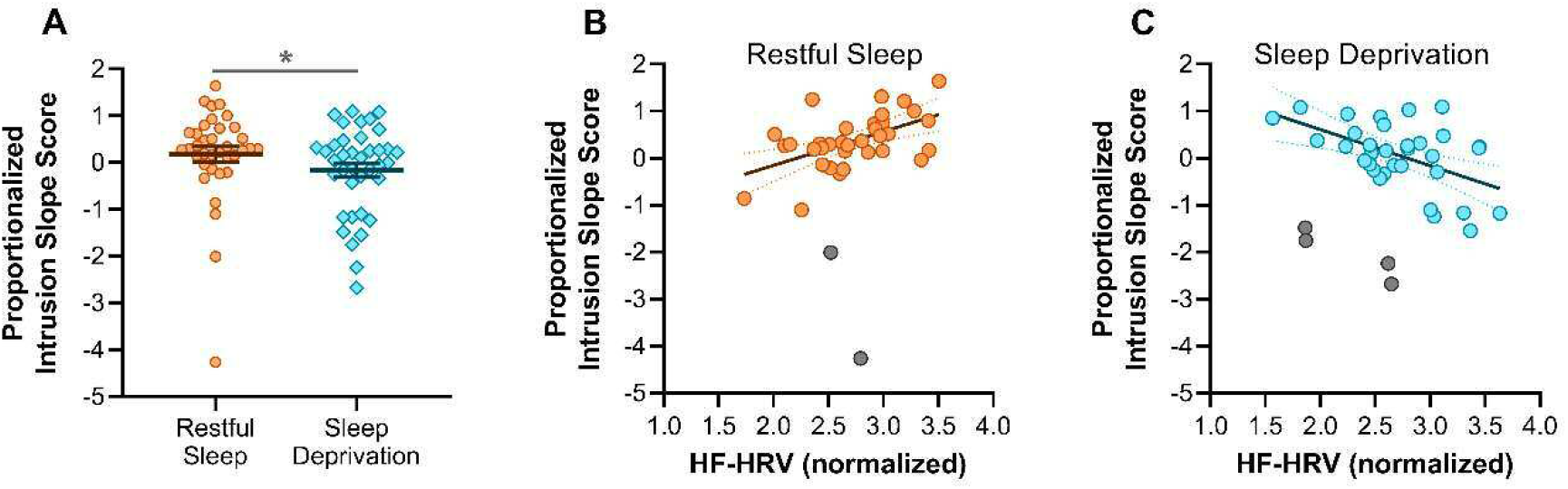
Adaptive memory suppression. (A) Intrusion slope scores were lower after sleep deprivation than restful sleep (a higher intrusion slope scores indicates a greater reduction in intrusions over trials). (B) In the restful sleep group, high-frequency heart rate variability (HF-HRV) was positively correlated with intrusion slope scores. (C) In the sleep deprivation group, HF-HRV was negatively correlated with intrusion slope scores. * = p<.05.

**Table 1.** Intrusion proportion (%), separately for each group and valence category.

|  | *Trial block* |  |  |  |  |
| --- | --- | --- | --- | --- | --- |
|  | 1 | 2 | 3 | 4 | 5 |
| ***Sleep-deprived*** |  |  |  |  |  |
| *Negative* | 36.09 (3.48) | 34.38 (3.30) | 29.86 (3.45) | 30.97 (3.33) | 29.45 (3.26) |
| *Neutral* | 37.05 (2.98) | 34.56 (3.13) | 30.73 (3.17) | 30.16 (3.54) | 31.02 (3.32) |
| ***Sleep-rested*** |  |  |  |  |  |
| *Negative* | 43.58 (3.81) | 38.25 (3.87) | 32.12 (3.69) | 28.90 (3.42) | 32.47 (3.73) |
| *Neutral* | 47.22 (4.22) | 39.73 (4.23) | 33.31 (4.23) | 32.49 (4.23) | 28.82 (3.88) |

Intrusion proportions are shown as means with SEM in parentheses.

Inhibitory control over cognition has been linked to the high-frequency component (0.15-40 Hz) of heart rate variability (HF-HRV)^10,11^. To investigate how trait HF-HRV influenced the impact of sleep deprivation on mnemonic control, we recorded resting heart rate before the overnight interval. Interestingly, higher HF-HRV was associated with higher intrusion slope scores, but only among participants who had slept (r_skipped_=.54 [0.22, 0.77]; Fig. 2B). For sleep-deprived individuals, higher resting HF-HRV was associated with *lower* intrusion slope scores (r_skipped_=-.52 [-0.75, −0.22], Zou’s CI [0.66, 1.34]; Fig. 2C). This suggests that individuals with higher trait HF-HRV are more adept at suppressing unwanted memories, but are also more susceptible to the disruptive influences of sleep deprivation on memory control. Further analysis of high-frequency and low-frequency components of resting HRV is available in Supplementary Fig. 1.

Although the foregoing analyses of intrusion slope scores suggest that adaptive memory suppression is impaired by sleep deprivation, there was no overall difference in intrusion proportion scores between the sleep-deprived and sleep-rested participants (F(1,72)=0.59, p=.44, *η_p_^2^*<0.01), irrespective of the emotional valence of the target scenes (F(1,72)=0.04, p=.84, *η_p_^2^*<0.01). These seemingly divergent findings may have arisen from between-group differences in memory control ability at baseline. For example, in a mock TNT task on unrelated practice items that took place before the overnight interval (see Methods), participants in the sleep deprivation group reported fewer intrusions than those in the sleep-rested group, although the between-group difference across this limited number of trials did not reach statistical significance (sleep-rested group: M=48.61%, SEM=3.76%; sleep deprivation group: M=38.82%, SEM=3.42%; F(1,72)=3.73, p=.058, *η_p_^2^*=0.05). The same pattern emerged in the first block of the main TNT assessment phase (t(72)=1.92, p=.059, d=0.45; Table 1), suggesting that the sleep-deprived participants were naturally more effective memory suppressors than the sleep-rested participants. Importantly, however, because the intrusion-reducing effect of repeatedly suppressing memory retrieval was larger in participants who had obtained restful sleep, the between-group difference in intrusion proportion scores was near zero by the final trial block (t(72)=0.09, p=.93, d=0.02). Hence, a lack of sleep appears to disrupt the cumulative and adaptive benefits of memory suppression, leading to a wholesale increase in vulnerability to intrusive memories. Consistent with earlier work^3,10^, intrusion proportion scores were not generally affected by the emotional valence of the target scenes (F(1,72)=0.29, p=.59, *η_p_^2^*<0.01).

#### fMRl

Next, we examined the impact of sleep deprivation on the neural correlates of memory suppression. Downregulating memories in response to reminders has previously been shown to engage rDLPFC^3,13–15,37,38^. Replicating this finding, our rDLPFC region of interest (ROI), which was obtained from an independent meta-analytic conjunction analysis of domain-general inhibitory control^13^, was more strongly engaged during memory suppression than retrieval (*F*(1,66)=55.69, *p*<.001, *η_p_^2^*=0.46). Critically, rDLPFC engagement during memory suppression (vs retrieval) was significantly lower after sleep deprivation than after restful sleep (interaction: *F*(1,66)=5.68, *p*=.020, *η_p_^2^*=0.08; Fig. 3A), suggesting that sleep loss impaired prefrontal memory control. Sleep deprivation (vs restful sleep) also led to a general reduction in rDLPFC activity (main effect: *F*(1,66)=11.58, *p*=.001, *η_p_^2^*=0.15). The effects of memory suppression and sleep deprivation on rDLPFC activation were not influenced by scene valence (all p≥.35), in keeping with prior work^3,10^.

**Figure 3.**
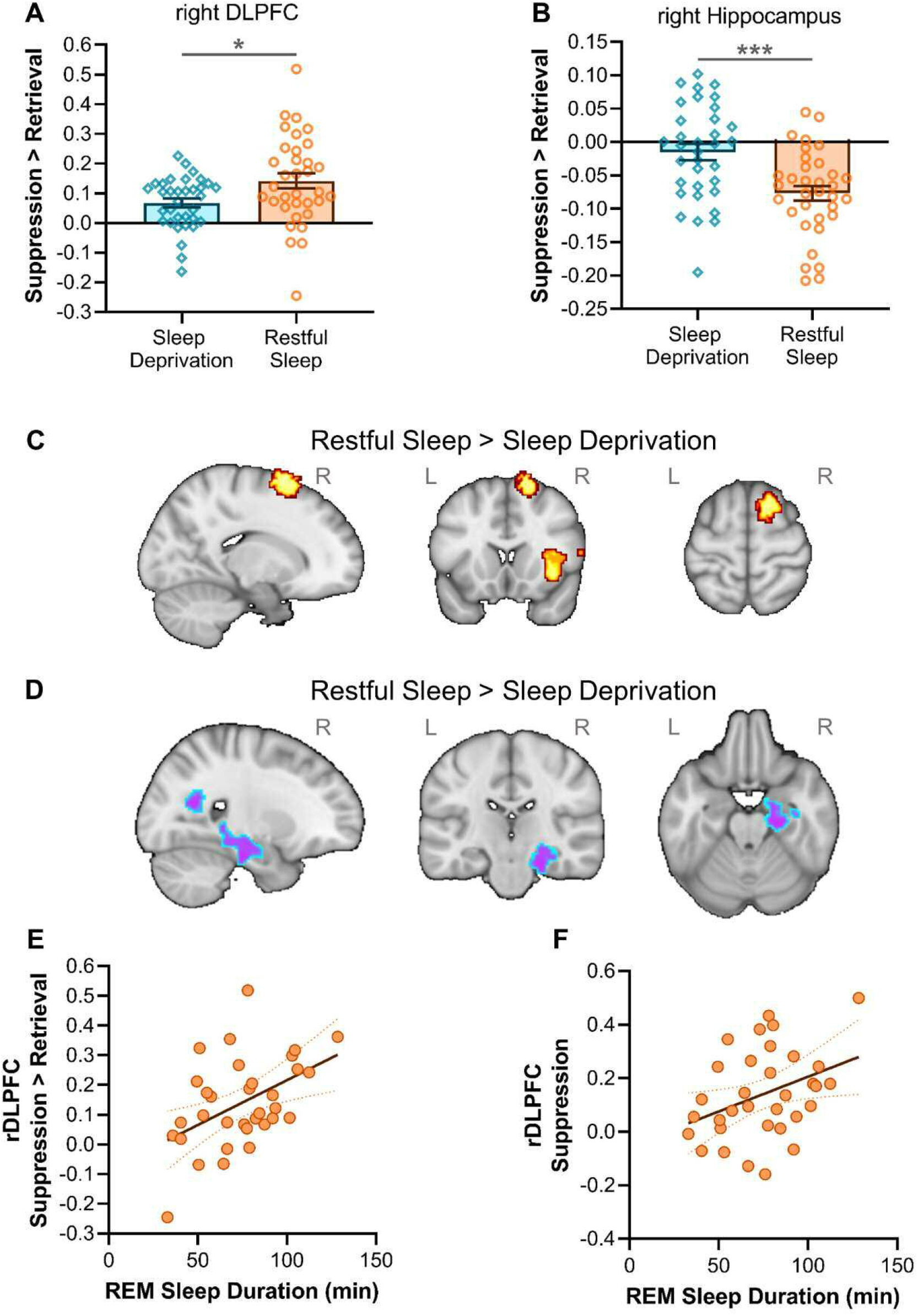
Functional brain responses during memory suppression (vs retrieval). Region of interest (ROI) analyses: (A) reduced engagement of right dorsolateral prefrontal cortex (rDLPFC) and (B) weaker disengagement of right hippocampus after sleep deprivation relative to restful sleep. Exploratory whole-brain analyses: (C) increased activation in right superior frontal gyrus and right insular cortex, and (D) decreased activation in right hippocampus after restful sleep (vs sleep deprivation). (E) Rapid eye movement (REM) sleep duration was correlated with right DLPFC engagement during memory suppression (vs retrieval) in the restful sleep group. (F) REM sleep duration was correlated with suppression-related activity in right DLPFC, whereas no such effect was observed for retrieval-related activity (not shown). * = p<.05; *** = p<.001, based on independent t-tests. R = right hemisphere; L = left hemisphere.

Memory suppression orchestrated by rDLPFC has been shown to target retrieval operations in hippocampus, especially in the right hemisphere^2,6,13,14,37,38^. In keeping with these prior findings, functional brain responses in our right hippocampus ROI were weaker during memory suppression than retrieval (*F*(1,66)=29.42, *p*<.001, *η_p_^2^*=0.31). Importantly, disengagement of right hippocampus during memory suppression (vs retrieval) was diminished after sleep deprivation relative to restful sleep (interaction: *F*(1,66)=17.02, *p*<.001, *η_p_^2^*=0.21; Fig. 3B), consistent with a failure to downregulate unwanted memories under conditions of sleep loss. No overall between-group difference was observed in right hippocampus (main effect: F(1,66)=3.12, p=.082, *η_p_^2^*=0.05), and the effects of memory suppression and sleep deprivation were not influenced by scene valence (all p≥.32).

We next investigated to what extent suppression-related activity in our ROIs was associated with behavioural expressions of memory control. We found that greater rDLPFC activation during suppression (vs retrieval) was associated with lower intrusion slope scores, but only among sleep-rested individuals (r_skipped_=-.40 [-0.63, - 0.11]). No such correlation was observed in sleep-deprived individuals (r_skipped_=-.21 [- 0.51, 0.16], Zou’s CI [-0.62, 0.26]). Similarly, rDLPFC activation during suppression (vs retrieval) was associated with higher intrusion proportion scores after restful sleep (r_skipped_=.42 [0.03, 0.69]) but not sleep deprivation (r_skipped_=.18 [-0.20, 0.46], Zou’s CI [- 0.21, 0.67]). Earlier work has shown that rDLPFC activity increases when unwanted memories enter into awareness and must be purged^2^. Individuals in the restful sleep group who experienced more mnemonic intrusions may have thus required stronger prefrontal engagement to reactively suppress unsolicited retrieval operations than those who had fewer intrusions. This dynamic prefrontal control mechanism had presumably failed after sleep deprivation, preventing participants from reactively purging intrusive memory content. Suppression-related activity in right hippocampus showed no relationship with intrusions in either the sleep-deprived (intrusion slope: r_skipped_=-.01 [-0.33, 0.33]; intrusion proportion: r_skipped_=.07 [-0.29, 0.35]) or sleep-rested individuals (intrusion slope: r_skipped_=-.08 [-0.46, 0.31]; intrusion proportion: r_skipped_=.01 [-0.33, 0.34]).

An exploratory whole-brain analysis contrasting memory suppression and retrieval (collapsed across negative and neutral scenes) showed that suppression-related activity in right superior frontal gyrus and right insular cortex was significantly higher in sleep-rested participants relative to sleep-deprived participants (Fig. 3C). Reciprocally, suppression-related activity in right hippocampus was significantly lower after restful sleep than sleep deprivation (Fig. 3D). Together with the findings from our ROI analyses, these data are consistent with a breakdown of prefrontal memory control as a consequence of sleep deprivation. Such a breakdown, theoretically, could set the stage for hippocampal hyperactivity and increased vulnerability to intrusive memories. See Table 2 for the full set of results from our whole-brain analysis.

**Table 2.** fMRI whole-brain analysis.

| *X Y Z (MNI)* | *No. voxels* | *Activation* | *Region* | *Z-Max* |
| --- | --- | --- | --- | --- |
| 14 10 72 | 462 | Increase | Superior frontal gyrus (right) | 5.00 |
| 36 14 -4 | 338 | Increase | Insular cortex (right) | 3.96 |
| 14 -52 18 | 4688 | Decrease | Precuneus (right) | 4.67 |
| 4 22 -2 | 1752 | Decrease | Subcallosal cortex (right) | 4.26 |
| 22 -22 -22 | 649 | Decrease | Parahippocampal gyrus (right) | 3.98 |
| -14 -86 2 | 350 | Decrease | Intracalcarine cortex (left) | 3.76 |

Activation changes associated with the contrast suppression>retrieval (collapsed across negative and neutral scenes) at the group level (restful sleep>sleep deprivation). X, Y, Z coordinates of the activation peak are shown in Montreal Neurological Institute (MNI) space. Cluster forming threshold $Z=3.1$ ; $p<.05$ (FWE corrected).

### REM sleep restores prefrontal memory control

Participants in the restful sleep group slept in our laboratory overnight so that we could examine the role of REM sleep in the overnight restoration of memory control. Interestingly, longer REM sleep duration was associated with greater rDLPFC engagement during memory suppression (vs retrieval; r_skipped_=.47 [0.17, 0.71]; Fig. 3E). Confirming that this effect was specific to memory suppression, REM sleep duration was significantly correlated with suppression-related responses (r_skipped_=.37 [0.07, 0.60]; Fig. 3F) but not retrieval-related responses in rDLPFC (r_skipped_=-.06 [-0.40, 0.29]), and these correlation coefficients were significantly different (Zou’s CI [0.12, 0.71]). REM sleep duration was not significantly correlated with activity in right hippocampus during memory suppression (vs retrieval; r_skipped_=.36 [-0.06, 0.71]). No significant correlation was observed between time spent in any non-REM sleep stage (N1, N2 or N3) and activity in rDLPFC or right hippocampus during memory suppression (vs retrieval). These findings point to a role for REM sleep in restoring prefrontal control mechanisms underpinning the ability to prevent unwanted memories from entering conscious thought.

### Sleep deprivation disrupts adaptive network segregation

So that we could assess the impacts of sleep deprivation on the brain’s intrinsic connectivity profile (i.e., in the absence of external task demands), participants underwent a resting-state fMRI scan after completing the TNT assessment phase. Sleep loss has been shown to disrupt functional decoupling between the DMN and CCN^23,28^, which support internally focused mental processing and externally directed cognitive control, respectively. In keeping with these earlier findings, DMN seed connectivity was significantly increased in a number of CCN areas—including bilateral middle frontal gyrus, inferior temporal gyrus, and supramarginal gyrus—after sleep deprivation relative to restful sleep (Fig. 4A). This suggests that sleep loss prevents the DMN from remaining functionally distinct from normally dissociable brain networks, giving rise to failures of cognitive control (see Supplementary Table 1 for the full set of clusters from our resting-state analyses).

**Figure 4.**
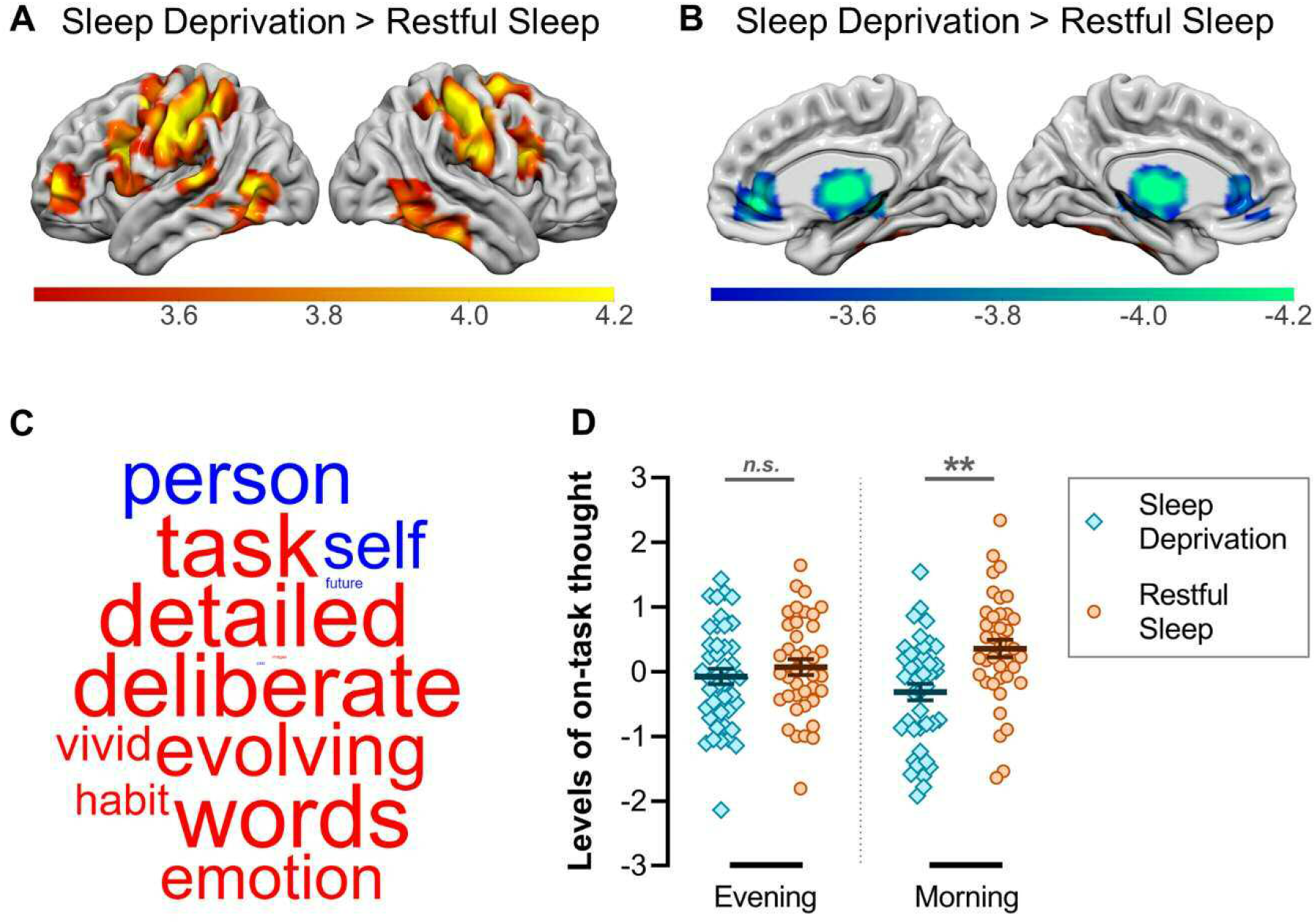
Resting-state functional connectivity and self-generated patterns of thought. (A) Sleep deprivation (vs restful sleep) increased functional connectivity between the default mode network (DMN) and several areas of the cognitive control network (CCN). (B) Sleep deprivation (vs restful sleep) decreased functional connectivity between the DMN and thalamus bilaterally. (C) The application of principal components analysis (PCA) to multidimensional experience sampling data identifies latent patterns of thought by grouping thought probes that capture shared variance. The first component identified from the PCA corresponded to a pattern of *on-task* thinking. The loadings on this component are presented as a word cloud. The colour of a word describes the direction of the relationship (red: positive, blue: negative) and the size of a word reflects the magnitude of the loading. (D) Patterns of on-task thinking were significantly reduced after sleep deprivation relative to restful sleep (no such difference emerged in the prior evening session). ** = p<.01; *n.s.* = non-significant.

Aberrant ascending input from thalamus to cortical regions encompassing the DMN and CCN has been implicated in the collapse of brain network integrity after sleep loss^29^. Consistent with this finding, sleep deprivation (vs restful sleep) significantly reduced DMN seed connectivity in bilateral thalamus (Fig. 4B). However, for the CCN seed, no differences in thalamic connectivity were observed between the sleep-deprived and sleep-rested participants. Disruptions to thalamocortical connectivity arising from sleep deprivation thus appear to predominate in the DMN and, in doing so, may contribute to a breakdown of adaptive functional segregation between the DMN and brain networks underpinning externally focused cognition.

### Sleep deprivation reduces deliberate patterns of thought

Finally, we examined how sleep deprivation affects the content of ongoing thoughts under conditions of high and low cognitive demand. In the evening before sleep deprivation or restful sleep, and again in the morning, participants completed a task that varied in its requirements for external attention (Fig. 1B). The task alternated between a condition with minimal attentional demands (0 back) and a higher demand condition in which task-relevant information had to be maintained in working memory (1 back). At intermittent intervals, participants were asked to describe the contents of their ongoing thoughts using multidimensional experience sampling (MDES)—an established thought sampling technique that is sensitive to task contexts and daily activities^39–41^. For this, they rated their recent experience along a series of thirteen thought probes, including whether or not their thoughts were focused on the task they were performing, were deliberate or spontaneous, or concerned past or future events (see Supplementary Table 2 for the full set of probes).

We applied principal components analysis to the MDES data for the purpose of identifying latent patterns of thought^39–41^. The first component identified in this manner captured an *on-task* dimension corresponding to deliberate and detailed thoughts about the task (Fig. 4C). Participants engaged in fewer of these on-task thoughts in the 0-back condition than in the relatively demanding 1-back condition (*F*(1,80)=30.68, *p*<.001, *η_p_^2^*=0.28), regardless of whether they were in the sleep deprivation or restful sleep group (*F*(1,80)=0.52, *p*=.47, *η_p_^2^*<0.01). Notably, while there was no group-level difference in on-task thinking at the evening session (before sleep deprivation or restful sleep, t=0.82, p=.41) one did emerge the following morning, with sleep-deprived participants exhibiting less on-task thinking than those who had slept (t=3.77, p=.001; interaction: *F*(1,80)=7.47, *p*=.008, *η_p_^2^*=0.09; Fig. 4D). This effect was independent of task demands (*F*(1,80)=0.69, *p*=.41, *η_p_^2^*<0.01). There was no overall difference in on-task thinking between the evening and morning sessions (*F*(1,80)=0.05, *p*=.83, *η_p_^2^*<0.01). These findings suggest that sleep loss impaired the ability to engage in deliberate, on-task thought, regardless of whether cognitive demands were high or low. See Supplementary Fig. 2 for analysis of the second component.

## Discussion

Our results demonstrate that sleep deprivation disrupts the neurocognitive mechanisms underpinning memory control. Relative to sleep-rested participants, sleep-deprived individuals were unable to properly engage rDLPFC during memory suppression, leading to a behavioural deficit in the ability to suppress unwanted memories over time. Among sleep-rested participants, longer REM sleep duration was associated with greater suppression-related activity in rDLPFC, suggesting that REM sleep supports the overnight restoration of prefrontal memory control. Consistent with prior work, sleep deprivation led to aberrant patterns of functional connectivity between brain networks underpinning internally and externally focused cognition^23,28,29^, as well as a tendency to engage in less deliberate, on-task thinking^30^.

It is now well established that rDLPFC downregulates hippocampal retrieval operations in service of memory control^2,3,13,14^. We previously showed that sleep deprivation leads to a marked reduction in people’s ability to suppress unwanted memories^10^, prompting our hypothesis that disruption to the prefrontal-hippocampal memory control network gives rise to memory suppression failures after prolonged wakefulness^16^. Our findings provide robust support for this hypothesis: sleep deprivation not only reduced rDLPFC engagement during memory suppression (as compared to restful sleep), but also weakened suppression-related activity in right hippocampus.

These findings add to a growing literature on the impacts of sleep deprivation on prefrontal control and underscore the importance of such findings for our understanding of mental health conditions that co-occur with chronic sleep disturbances. For example, sleep loss impairs prefrontal control mechanisms that resolve conflict between habitual and goal-directed behaviour^42^, providing mechanistic insight into the link between sleep disturbance and relapse in individuals with addictive disorders^43^. Likewise, trait-anxious individuals and patients with anxiety disorders show reduced prefrontal activation and impaired functional coupling between prefrontal cortex and amygdala during threat-related information processing^44,45^, a pattern that also emerges in healthy individuals deprived of sleep^20,46^. Given that memories play a central role in our affective perception of the external world^47^, memory control failures may go a long way towards explaining the relationship between sleep loss and emotional dysregulation.

Along similar lines, we found that the magnitude of rDLPFC engagement during memory suppression was correlated with the amount of time that sleep-rested participants spent in REM sleep. Disturbances of REM sleep (e.g., reduced REM sleep latency) are commonplace in psychiatric disorders associated with intrusive and unwanted thoughts, including depression, anxiety and post-traumatic stress disorder^32,34,36^. Furthermore, in healthy individuals, REM sleep has been implicated in affect regulation during exposure to emotionally aversive stimuli^48,49^. Taken together, these and the current findings raise the intriguing possibility that REM sleep supports the restoration of memory and emotional control processes mediated by prefrontal cortex. Future work can address this possibility by directly intervening with REM sleep (e.g., via non-invasive auditory brain stimulation^50^) and assessing its causal impacts on memory and affective suppression.

Our exploratory whole-brain analysis showed that sleep deprivation reduced activity in right insular cortex during memory suppression. This likely reflects an impact of sleep loss on the brain’s ability to switch between activation and deactivation of large-scale networks. Indeed, right insular cortex plays a major role in switching between distinct brain networks across task paradigms and stimulus modalities^51^, as is the case in our Think/No-Think protocol, where participants must rapidly shift between conditions of memory retrieval (Think trials) and suppression (No-Think trials). In keeping with this idea, functional networks encompassing right insular cortex show reduced activity when performing attention-demanding tasks after sleep deprivation^52^. Relatedly, monitoring one’s own internal state (as is required when repeatedly switching between conditions with differing cognitive demands) may benefit from interoceptive processing, which recruits a network including insular cortex^53^, and is impaired by poor quality sleep^54^.

The findings from our resting-state fMRI data were consistent with prior evidence that sleep deprivation leads to aberrant patterns of functional connectivity between whole-brain functional networks supporting internally and externally focused cognition^23,28^. In particular, we observed an increase in connectivity between the DMN and multiple areas of the CCN in sleep-deprived individuals, as compared to sleep-rested individuals. Moreover, DMN connectivity with bilateral thalamus was significantly reduced after sleep deprivation relative to restful sleep, in keeping with earlier work^29^.

High-frequency heart rate variability (HF-HRV) is linked to superior executive functioning, including memory and emotional control^10,12^. Consistent with this earlier work, our data revealed a significant correlation between HF-HRV at rest and memory suppression success over time. Importantly, whereas previous studies have linked HF-HRV to suppression-induced forgetting (i.e., lower recall for suppressed relative to baseline word pairs)^11^, this is the first study to demonstrate that resting cardiac activity can reliably predict a person’s ability to directly suppress unwanted memories. This relationship was only observed in individuals who had obtained a night of restful sleep; for sleep-deprived participants, the opposite effect was observed: that is, higher resting HF-HRV was associated with *lower* memory suppression success. This provides the first indication that people who are inherently resilient to intrusive memories might also be the most vulnerable to the effects of sleep deprivation.

Our multidimensional experience sampling (MDES) protocol allowed us to examine how sleep deprivation affects the content of self-generated thoughts. Whereas the sleep-deprived and sleep-rested groups showed no difference in deliberate and detailed patterns of thinking prior to the overnight delay (based on an *on-task* thought dimension derived from principal components analysis of the MDES data), a group-level difference did emerge the following morning: sleep-deprived individuals reported fewer deliberate, on-task thoughts than sleep-rested participants, which is in keeping with prior work using self-reported categorisations of on-task and off-task thought^30^. The effect of sleep loss on on-task thinking was unaffected by task demands, suggesting that an absence of sleep leads to significant changes in the content of endogenous thoughts that are impervious to external contexts. Interestingly, previous work has shown that sleep-deprived individuals tend to rely on habits at the expense of goal-directed control during decision making^42^. A generalised reduction in on-task thinking after sleep deprivation might thus reflect a deficit in the ability to align cognition with goals of the moment, which could in itself give rise to intrusive and unwanted thoughts.

In conclusion, our findings demonstrate that sleep deprivation leads to widespread disruption of the brain networks supporting adaptive control. An absence of sleep impaired rDLPFC engagement during memory suppression, increased suppression-related activity in hippocampus and disrupted resting-state connectivity between brain regions supporting internally (i.e., DMN) and externally focused cognition (CCN). Among participants who obtained restful sleep, engagement of rDLPFC during memory suppression was correlated with time spent in REM sleep, suggesting that REM sleep might play a central role in the overnight restoration of prefrontal memory control mechanisms. The functional impairments arising from sleep deprivation were associated with a behavioural deficit in suppressing unwanted memories over time and coincided with a deterioration of deliberate, on-task thinking. Taken together, our findings highlight the critical role of sleep in maintaining control over both our memories and ongoing thoughts.

## Methods

### Participants

Eighty-seven healthy adults completed the experiment. Participants were right-handed, native English speakers with no history of neurological, psychiatric, attention, or sleep disorders. According to self-report, they typically awoke by 8am after at least 6 hours of sleep. Following standard procedures in our laboratory^55–59^, participants were asked to abstain from alcohol and caffeine for 24 h prior to the experiment. Written informed consent was obtained from all participants in line with the requirements of the Research Ethics Committee of the York Neuroimaging Centre at the University of York, who approved the study. Participants were compensated with £80 or experimental participation credit (University of York BSc Psychology students only).

Two participants were excluded from all analyses due to lack of appropriate engagement with the study protocol (e.g., persistently failing to follow instructions). The final sample included 85 participants, who completed either the sleep deprivation condition (n=43; 18 males, mean ± SD age, 19.58 ± 1.72 years) or restful sleep condition (n=42; 12 males, mean ± SD age, 20.33 ± 2.43 years).

### Procedure

Two sessions (evening and morning) were separated by a night of sleep deprivation or restful sleep (see Fig. 1A). Participants collected an actigraphy wristwatch (Philips Respironics, Murrysville, PA) at 9am on the day of the evening session so we could ensure they had not napped during the day (confirmed via subsequent analysis of actigraphy data). Hence, by the time of the morning session, participants in the sleep deprivation group had been awake for ∼24 hours.

#### Evening

T1 structural MRI scans were collected in the evening (∼6pm) to ensure that participants were comfortable in the MRI environment before committing to the full experiment. After participants exited the scanner, an 8-min resting electrocardiography (ECG) recording was obtained for the purpose of calculating heart rate variability (HRV). Participants were instructed to sit with their hands on their lap, relax, and breathe normally throughout.

We then administered the multidimensional experience sampling (MDES) task (Fig. 1B). Experience was sampled in a task that included non-target and target trials and switched between conditions of 0-back and 1-back for the purpose of manipulating working memory demands. Non-target trials were identical in the 0-back and 1-back conditions and consisted of two different shapes (circles, squares, or triangles) separated by a centre line (jittered duration from 500–1500 ms). The colour of the centre line indicated to the participant which condition they were in (blue = 0-back; red = 1-back). Non-target trials did not require a behavioural response from participants and were presented in runs of 2-5 trials, after which a target trial or MDES probe was presented. On target trials, participants were required to indicate the location of a particular shape (circle, square, or triangle; jittered duration from 3500–5000 ms). The cognitive demand required to fulfil this instruction depended on whether participants were in the 0-back or 1-back condition. In the 0-back (non-demanding) condition, for target trials, two different shapes were presented on either side of the centre line (as in the non-target trials), and an additional shape appeared in the middle of the centre line. Participants were required to indicate (via button press) which of the lateral shapes matched the central shape. Hence, in the 0-back condition, non-target trials did not require continuous monitoring and participants could make perceptually guided decisions. In the 1-back condition, target trials also involved a shape appearing in the middle of the centre line, but with a question mark on either side (instead of shapes). Participants were required to indicate which of the two lateral shapes from the *previous* trial matched the present central shape. Target trial decisions in the 1-back condition were therefore guided by working memory, meaning that the participants were required to continuously monitor the non-target trials. All participants completed the 0-back and 1-back conditions once (order counterbalanced), with two target trials per condition. Patterns of ongoing thought were measured using MDES probes, which occurred instead of target trials on a quasi-random basis. For the MDES probes, participants were asked to what extent their thoughts were focused on the task, followed by 12 randomly shuffled questions about the content of their thoughts (see Supplementary Table 2). The questions were administered four times (twice in each of the 0-back and 1-back conditions) and participants made their responses on a sliding scale ranging from 1 to 10.

Affect ratings were then collected for 68 scenes. Each scene was presented for 6500 ms and participants were instructed to focus their attention on the image for the entire time. They were then shown a 9-point pictorial rating scale that ranged from a frowning face on the far left (depicting feelings of extreme sadness or displeasure) to a smiling face on the far right (depicting feelings of extreme happiness or pleasure).

Participants were given 15000 ms to provide their affect rating (1 to 9) via button press, but were asked to respond quickly and spontaneously. Trials terminated after an affect rating had been provided or the 15000 ms time limit expired, after which a fixation cross appeared for 500 ms, 1000 ms, 1500 ms, or 2000 ms. Affect rating data is available in Supplementary Table 3.

Participants then completed the Think/No-Think learning phase, which involved encoding pairwise associations between faces and scenes (Fig. 1C). On each trial, participants were shown a face and scene together for 6000 ms and instructed to form a rich connection between them. Faces were always emotionally neutral whereas half the scenes were negative and the other half were neutral. Participants then completed a reinforcement phase where they were shown each of the faces in isolation for up to 4000 ms and indicated via button press whether or not they could visualize the corresponding scene. When participants indicated that they could visualize the scene, they were shown the correct scene alongside two additional scenes from the learning phase that were not paired with the face and asked to select the correct image. If participants selected the correct scene, the face-scene pair was dropped from the reinforcement phase. If they failed to select the correct scene within 5000 ms or they had indicated that they could not visualize the scene associated with the face, the face-scene pair was retained in the reinforcement phase and they were tested on it again later in the same phase. Regardless of their response, participants were always shown the correct face-scene pairing for 3500 ms at the end of the trial to reinforce their knowledge of the pairs. The reinforcement phase continued until participants had correctly identified the target image associated with each face cue. Participants then completed the entire reinforcement phase again. This ‘overtraining’ procedure was used to ensure that participants would find it difficult to prevent the scenes from automatically intruding into consciousness when presented with the face cues during the TNT assessment phase. Forty-eight face-scene pairs (24 negative, 24 neutral) were presented at the learning phase and used in the main TNT assessment. Twelve additional face-scene pairs (6 negative, 6 neutral) served as fillers that were also used in the mock TNT assessment.

The mock version of the TNT assessment phase was then administered (outside of the MRI scanner) to enable participants to practice engaging in memory suppression. On each trial, a face cue was presented in isolation inside a green frame (*Think* trial) or red frame (*No-Think* trial). For green-framed faces, participants were instructed to visualize—in as much detail as possible—the scene associated with the face for the entire 3000 ms that it was presented. For red-framed faces, participants were instructed to focus their attention on the face for the entire 3000 ms, but simultaneously prevent the associated scene from coming to mind. Participants were instructed to accomplish this by making their mind go blank, rather than by generating diversionary thoughts such as alternative images, thoughts, or ideas^3,8,10^. If the scene came to mind automatically, participants were asked to actively push the scene out of mind. At the offset of each face cue, participants reported whether the scene paired with the preceding face had entered conscious awareness during the trial by pressing (with their right hand) a button corresponding to one of three options: *never*, *briefly*, or *often*. Participants were instructed to provide a response of *never* if the scene never entered awareness at all during the trial. Conversely, participants were instructed to respond with *briefly* if the scene briefly entered conscious awareness at any time during the trial, or with *often* if the scene entered awareness several times or for a period longer than what one would consider ‘brief’. These intrusion ratings were collected to determine how competent participants were at suppressing scenes associated with faces for No-Think trials. Although participants had up to 4000 ms to make this rating, they were instructed to respond quickly, without dwelling on their decision. Participants moved immediately onto the next trial after providing their rating (jittered fixation of 500 ms, 2000 ms, 2500 ms, 4000 ms, or 6500 ms). The mock TNT assessment included only faces from the 12 face-scene pairs used as fillers at learning, and these were equally divided between conditions (6 Think, 6 No-Think; each 3 negative, 3 neutral).

#### Overnight

The restful sleep group were fitted with electrodes for sleep polysomnography. Lights were turned out at approximately 10pm and participants were awoken at 6am, providing an 8 h sleep opportunity. The sleep deprivation group remained awake across the entire night either in a university seminar room under the supervision of an experimenter (n=23) or at home (n=20). The reason why some participants stayed awake at university and others at home was due to UK Government social distancing guidelines during the SARS-CoV-2 pandemic, which came into force part-way through the experiment. To ensure that participants who stayed awake at home adhered to the sleep deprivation protocol, they were asked to send an SMS message to the experimenter every 30 min throughout the night. This requirement was fulfilled by all 20 participants. Wristwatch actigraphy also confirmed that these participants remained awake throughout the night. During the overnight phase, the sleep-deprived participants were permitted to read, use personal computers or other devices, watch TV, or play games. They were also permitted to eat and drink at any time, but caffeine was prohibited.

#### Morning

A second ECG recording was obtained from each participant under the same conditions as the preceding session. Participants then entered the MRI scanner. We first administered a memory refresher phase where each of the face-scene pairs were presented for 1.5 s, allowing participants to reinforce their knowledge of the pairs (no scanning was performed at this time). The TNT assessment phase proper was then administered in 5 blocks while participants were undergoing fMRI. This followed the same procedures as the mock TNT assessment, but included faces from the main 48 face-scene pairs presented at learning (faces from the filler pairs used in the mock TNT assessment were not included here) and had a jittered fixation of 500 ms – 9000 ms between trials. Each block lasted approximately 8 min and included two repetitions of 16 Think (8 negative, 8 neutral) and 16 No-Think (8 negative, 8 neutral) items presented in pseudorandom order (with the two repetitions of each item appearing at least three trials away from one another). Participants therefore completed a total of 320 trials (32 trials x 2 conditions x 5 blocks). The remaining 16 scenes that were included in the TNT learning phase did not appear in the TNT assessment phase. These baseline images were used in the affect rating task to index generalized changes in emotional reactivity that could be compared to changes arising from mnemonic suppression (see Supplementary Table 3). A B0 fieldmap was acquired between the second and third blocks of the TNT assessment.

Participants then underwent a 9-min resting-state fMRI scan and were instructed to focus on a fixation cross in the centre of the screen throughout. After participants exited the scanner, we collected another round of affect ratings and administered the MDES paradigm again, following identical procedures to the evening session. Finally, participants completed a post-experiment questionnaire, which probed how closely they had followed the TNT task instructions^60^.

### Stimuli

Forty-eight emotionally neutral face images (24 male, 24 female) served as cues in the TNT task, and an equal number of scene images (24 negative, 24 neutral; Lang et al., 2005) served as targets. Face-scene pairs were created by randomly assigning each face cue to a target scene. Three lists of 16 pairs (8 negative; 8 neutral) were created from the 48 pairs to allow three within-subjects TNT conditions (Think, No-Think, and Baseline). A second version of each of the three lists was also created by assigning a new face cue to each target scene within a given list. The assignment of face-scene pairs to TNT conditions was counterbalanced across participants, as was the version of face-scene pairings within the TNT condition. Twelve additional face-scene pairs (6 negative, 6 neutral) were created for use as fillers. The same 60 scenes (48 experimental + 12 fillers) were used for the affect rating task, which featured a further 8 filler scenes (4 negative, 4 neutral).

### Software

The TNT and MDES tasks were run in Presentation version 20.3 (Neurobehavioral Systems, Albany, CA) and PsychoPy2^62^, respectively. Tasks administered outside of the MRI scanner were run on a desktop PC with flat-screen monitor or laptop, and responses were collected using the keyboard.

### MRI data acquisition

MRI scans were performed at York Neuroimaging Centre, University of York, UK, with a Siemens MAGNETOM Prisma 3T scanner. All scans were acquired with a 64-channel head coil, with whole brain coverage.

#### T1 structural scan

A high resolution T1-weighted structural scan was acquired with a 3D magnetization-prepared rapid gradient-echo (MPRAGE) sequence (TR=2300 ms, TE=2.26 ms, flip angle=8°, field of view = 256mm, isotropic voxel size=1mm, 176 slices, slice thickness=1 mm, anterior-to-posterior phase encoding direction, in-plane acceleration factor=2 [GRAPPA]).

#### fMRI

Event-related changes in blood oxygen level-dependent (BOLD) signal were acquired with a T2*-weighted gradient echo-planar imaging (EPI) sequence (TR=2000 ms, TE=30 ms, flip angle=75°, field of view = 240mm, isotropic voxel size=3mm, 70 slices, slice thickness=3mm, interleaved slice acquisition, anterior-to-posterior phase encoding direction, in-plane acceleration factor=2 [GRAPPA]). Five event-related fMRI scans were acquired per participant (corresponding to 5 blocks of the TNT assessment phase; referred to hereafter as TNT scans). Each scan was preceded by dummy volumes to allow for steady-state magnetizations to become established (at which time the scanner’s trigger signal initiated the task in synchrony with the acquisition of the first fMRI volume proper). Stimuli were projected onto a screen at the rear of the magnet bore with a PROPixx DLP LED Projector (VPixx Technologies Inc) and comfortably viewed with an angled mirror attached to the head coil. Participant responses were captured with a USB button response box placed in their right hand.

A 9-min resting-state fMRI scan was acquired immediately after participants completed the TNT task with a gradient EPI sequence (TR=2000 ms, TE=30 ms, flip angle=80°, field of view = 240mm, isotropic voxel size=3mm, 70 slices, slice thickness=3mm, interleaved slice acquisition, anterior-to-posterior phase encoding direction, in-plane acceleration factor=2 [GRAPPA]).

#### B0 fieldmap

We acquired a B0 fieldmap to correct for magnetic field inhomogeneities in the fMRI data (TR=850 ms, TE1=4.92 ms, TE2=7.38 ms, flip angle=60°, field of view = 240mm, isotropic voxel size=3mm, 72 slices, slice thickness=3 mm, interleaved slice acquisition, right-to-left phase encoding direction). The B0 fieldmap was acquired between the second and third TNT block. If participants exited the scanner for a break between scans (n=3), a second fieldmap was acquired after they re-entered the scanner. This second B0 Fieldmap was used to correct for magnetic field inhomogeneities in the remaining scans.

### Heart rate variability

Electrocardiography (ECG) was recorded using a BIOPAC MP36R data acquisition system and AcqKnowledge version 4.4.1. Three BIOPAC EL503 ECG electrodes were attached to the midline of the left and right clavicle and the lower left rib. ECG was recorded continuously at a sampling rate of 2 KHz for 8 min.

### Polysomnography

Sleep was recorded using an Embla N7000 polysomnography (PSG) system (Embla Systems, Broomfield, CO, USA) and RemLogic version 3.4. The scalp was cleaned with NuPrep exfoliating agent, before gold-plated electrodes were attached using SAC2 electrode cream. Electroencephalography (EEG) electrodes were attached at eight standard locations according to the international 10-20 system (Homan, Herman, & Purdy, 1987): F3, F4, C3, C4, P3, P4, O1, and O2, each referenced to the contralateral mastoid (A1 or A2). Left and right electrooculogram, left, right and upper electromyogram, and a ground electrode (forehead) were also attached. All electrodes were verified to have a connection impedance of < 5kΩ and were sampled at a rate of 200 Hz.

To calculate the time spent in each stage of sleep, the PSG recordings were partitioned into 30 s epochs and scored, using RemLogic software, as wakefulness, N1, N2, N3, or rapid eye movement (REM) sleep in accordance with the criteria of the American Academy of Sleep Medicine^63^.

### Data analysis

Data were analyzed using JASP 0.13.1.0 (JASP Team, Amsterdam, The Netherlands), unless specified otherwise.

#### Behaviour

Although intrusion reports for each trial of the TNT assessment phase were obtained on a 3-point scale (*never*, *briefly*, *often*), participants rarely gave ratings of *often* for No-Think trials (mean ± SEM, 2.47 ± 0.49 %). For simplicity, we therefore followed our prior work^10^ by combining the *briefly* and *often* responses, rending the judgement binary. Intrusion proportion scores were robustly greater for Think trials (mean ± SEM, 93.25 ± 0.92 %) as compared to No-Think trials (mean ± SEM, 34.01% ± 2.11 %; t(73)=25.22, p<.001, d=2.93), demonstrating that participants were generally effective at voluntarily preventing unwanted memories from entering awareness. Because our research questions concern involuntary memory intrusions, our analyses focus exclusively on No-Think trials.

Intrusion proportion scores were applied to a mixed measures ANOVA with factors Group (Sleep Deprivation, Restful Sleep), Emotion (Negative, Neutral), and Task Block (1, 2, 3, 4, 5). Mauchly’s test of sphericity indicated that for some effects the assumption of sphericity was violated. In these cases, Greenhouse-Geisser correction was applied. To directly quantify adaptive memory suppression (i.e., the rate at which intrusions decreased over time), we computed an intrusion slope score for each participant. Intrusion slope scores were calculated by taking the slope of the linear regression line through intrusion proportion scores (averaged across scene valence categories) across TNT trial blocks. This value was divided by the participant’s intrusion proportion score in the first block to account for the fact that initial intrusion rates varied and participants with more initial intrusions had a greater margin for reducing their intrusion frequency. We then multiplied the values by –1 to render the (primarily) negative scores positive. Increasing positive scores thus reflect increasing levels of competency at down-regulating the frequency of intrusions over time. This measure was z normalized within the participant’s stimuli counterbalancing group^4,5,10^, which allowed us to quantify intrusion slope scores with respect to a group of participants who attempted to suppress/retrieve precisely the same scenes in the TNT task. Shapiro-Wilk tests indicated that intrusion slope scores were not normally distributed in either group (both *p* < .05), so non-parametric Mann-Whitney U tests were used to compare intrusion slope scores between the sleep-deprived and sleep-rested groups. N=11 participants were excluded from the TNT behavioural analyses due to: revealing in the post-experiment questionnaire that they had engaged in unsolicited behaviours during the TNT task (e.g., intentionally thinking about scenes associated with red-framed faces; n=8)^60^, indicating that they had not understood the task instructions (n=2), or data loss due to a technical fault (n=1).

Skipped Pearson’s correlations (MATLAB toolbox: Robust Correlation)^64^ were used to investigate linear relationships between variables (e.g., functional brain activity and time spent in REM sleep). Skipped correlations detect and ignore outliers by considering the overall structure of the data, providing accurate false positive control without loss of power. We compared the skipped correlations between groups using Zou’s confidence intervals (CI; R package: cocor)^65^. Skipped correlations were interpreted as significantly different if Zou’s confidence interval did not contain zero^65^. N=2 participants were excluded from correlational analyses involving PSG data due to EEG electrodes becoming detached during the night (preventing accurate calculation of time spent in each sleep stage).

#### MDES

Principle components analysis (PCA) was carried out in SPSS version 27.0.1.0. The scores from the 13 experience sampling questions were entered into a PCA to describe the underlying structure of the participants’ responses. Because the 13 thought probes were administered twice during each of the 0-back and 1-back blocks, the two responses to each probe were averaged across iterations. Following our prior work^41^, we concatenated the 13 responses of each participant in each session (evening/morning) and task (0-back/1-back) into a single matrix and carried out a PCA with varimax rotation. Two components were selected based on the inflection point in the scree plot (see Supplementary Fig. 2), cumulatively explaining 42.77% of the total variance. The first component corresponded to a pattern of on-task thought, based on its question loadings. Scores for the two components were extracted for each participant and entered into a mixed-measures ANOVA with factors Task (0-back, 1-back), Session (Evening, Morning) and Group (Sleep Deprivation, Restful Sleep). Significant interactions were interrogated using Holm-Bonferroni post-hoc comparisons. N=3 participants were excluded from the MDES analysis due to their data not being acquired in both sessions (N=2) or providing very high ratings to an unrealistic number of thought probes (N=1; mean response of 9.3/10, which was >4SD above the group mean).

#### fMRI

Event-related fMRI data were analysed using the FMRIB Software Library (FSL version 5.0; https://fsl.fmrib.ox.ac.uk/fsl/fslwiki/FEAT/). T1-weighted structural brain images were extracted using the Brain Extraction Tool (BET) and the functional data were preprocessed and analysed using the fMRI Expert Analysis Tool (FEAT). Individual TNT scans were subjected to motion correction using a six-parameter rigid body transformation, B0 unwarping and slice-timing correction. TNT scans were co-registered with the relevant structural image using Boundary-Based Registration^66^ and were spatially normalized to the Montreal Neurological Institute (MNI-152) canonical brain. Functional images were spatially smoothed using a Gaussian kernel of FWHM 8mm and underwent high-pass temporal filtering (Gaussian-weighted, 100 s).

A general linear model (GLM)^67^ was set up for each participant and TNT scan. This included four explanatory variables (EVs) to model retrieval of negative scenes (EV1: Think/Negative), retrieval of neutral scenes (EV2: Think/Neutral), suppression of negative scenes (EV3: No-Think/Negative) and suppression of neutral scenes (EV4: No-Think/Neutral). All of the EVs were convolved with a canonical haemodynamic response function, and their temporal derivatives were also added to the model. Six movement parameters, acquired during motion correction, were included as non-convolved nuisance regressors. Contrast images were obtained for each effect of interest (Think/Negative, Think/Neutral, No-Think/Negative, No-Think/Neutral). For each participant, the parameter images corresponding to each TNT scan were taken forward to a higher-level, fixed-effects model to average across activity within individuals.

Our region-of-interest (ROI) analyses focused on right dorsolateral prefrontal cortex (rDLPFC) and right hippocampus, as these regions have been heavily implicated in memory suppression. For the rDLPFC ROI, we constructed a 5 mm sphere centered at MNI coordinates 35, 45, 24, which were obtained from an independent meta-analytic conjunction analysis of domain-general inhibitory control^13^. The ROI for right hippocampus was taken from the Harvard-Oxford subcortical atlas (http://fsl.fmrib.ox.ac.uk/fsl/fslwiki/Atlases) and was restricted to voxels with ≥50% probability of belonging to right hippocampus. Individual parameter estimates were extracted from the higher-level fixed-effects analyses (averaging across TNT scans in each participant) and averaged in each ROI (using the featquery tool). The ROIs were then separately analysed using a mixed-measures ANOVA with factors Group (Sleep Deprivation, Restful Sleep), Memory Process (Retrieval, Suppression) and Emotion (Negative, Neutral). Given no effects of Emotion emerged in either ROI, we collapsed across valence categories in subsequent Suppression>Retrieval analyses.

A complementary whole-brain analysis was conducted in FSL to permit further investigation of how sleep deprivation affects the neural correlates of memory suppression. Because our ROI analyses revealed no significant effects of Emotion, we set up new GLMs for each participant and TNT scan with two EVs (Think and No-Think) that were both collapsed across negative and neutral scenes (the set up for this first-level analysis was otherwise identical to that reported above). A contrast was obtained for the effect of No-Think>Think, and the resulting contrast images for each TNT scan were averaged within participants using a higher-level fixed-effects analysis. Participant contrast images were then used as input for the group-level (mixed-effects) analysis. For this, we used FLAME, as implemented in FSL, with a cluster forming threshold of Z=3.1 and p<.05 family-wise error correction. N=17 participants were excluded from the event-related fMRI analyses due to: being excluded from the behavioural analysis (n=11; see *Data Analysis* for further details), poor co-registration between the EPI and structural images (n=2), or loss of ≥3 (out of 5) TNT scans due to excessive movement (relative displacement >3mm) and/or major artifact in the EPI data (n=4). For participants with excessive movement detected on ≤2 TNT scans (relative displacement >3mm; n=5), those volumes were excluded and higher-level analyses were conducted on the remaining data.

Resting-state fMRI data were analysed using the CONN functional connectivity toolbox version 21.a (https://web.conn-toolbox.org/)^68^, implemented with SPM version 12 (https://www.fil.ion.ucl.ac.uk/spm/) and MATLAB version 2019a. Preprocessing steps followed CONN’s default pipeline, which included motion correction using a six-parameter rigid body transformation, slice-time correction and the simultaneous segmentation and normalization of grey matter, white matter and cerebrospinal fluid within both the fMRI and T1-weighted structural data to the MNI-152 canonical brain. Next, we removed potential confounding effects from the BOLD signal using linear regression, including the six movement parameters acquired during motion correction and their 1^st^ and 2^nd^ order derivates, volumes with excessive movement (motion greater than 0.5mm and global signal changes larger than z=3), signal linear trend, and five principal components of the signal from white matter and cerebrospinal fluid (CompCor)^69^. A bandpass filter of 0.01-0.1 Hz was also applied to the data.

Seed-based connectivity maps were calculated for each participant with the default mode network and frontoparietal cognitive control network as ROIs (as derived from the Yeo 7-network parcellation)^70^. Nuisance regressors included movement parameters calculated during motion correction, volumes with excessive movement and framewise displacement. The participant connectivity maps were then used as input for the group-level analysis, which included age, gender and mean framewise displacement as mean-centered covariates. All connectivity results were thresholded using Gaussian Random Field theory, as implemented in CONN, with a p<0.001 (uncorrected, two-sided) voxel threshold and an FDR-corrected cluster threshold of p<0.05/7=0.007, to also correct for testing multiple seeds (Bonferroni correction). N=6 participants were excluded from the resting-state analysis due to: poor co-registration between the EPI and structural images (n=2), excessive movement (n=2), a major artifact in the structural image (n=1), or a neurological anomaly (n=1; note that this participant was also excluded from the behavioural and event-related fMRI analyses for indicating that they had misunderstood the TNT task instructions).

#### Heart rate variability

The first 2-min and last 1-min of each 8-min ECG recording were excluded and HRV was calculated using the remaining 5-min of data. R-peaks were automatically detected using AcqKnowledge’s QRS detection algorithm before being visually inspected for accuracy. Peaks that the algorithm missed or erroneously detected were manually inserted or deleted, respectively. The interbeat-interval time series was then imported to Kubios version 3.0.2. for analysis. Autoregressive estimates of low-frequency (0.04-0.15 ms2/Hz) and high-frequency (0.15-0.40 ms2/Hz) power were used to obtain frequency-domain-specific indices of HRV. In keeping with earlier work^11,71^, values of low-frequency HRV (LF-HRV) and high-frequency HRV (HF-HRV) were transformed logarithmically (base 10). N=1 participant was excluded from all HRV analyses because the ECG revealed signs of cardiac arrhythmia.

## Acknowledgements

This work was supported by Medical Research Council Career Development Award MR-P020208-1 to SAC.

