## Supplementary for "Memory control deficits in the sleep-deprived human brain"

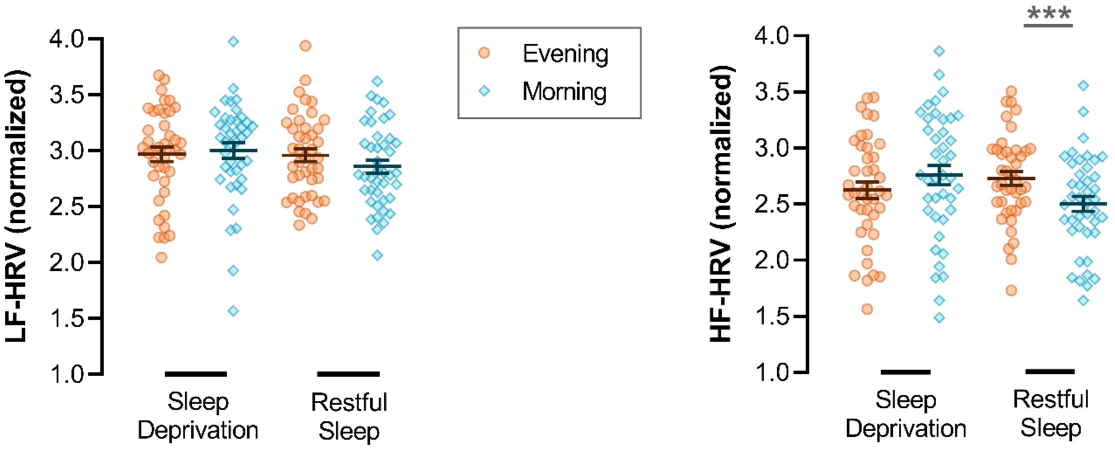


**Supplementary Figure 1.** Low-frequency and high-frequency heart rate variability (LF-HRV; HF-HRV). To investigate the impact of sleep loss on HRV, our indices of HF- and LF-HRV were applied to mixed 2 (Group: Sleep Deprivation, Restful Sleep) x 2 (Session: Evening, Morning) ANOVAs. Post-hoc comparisons were conducted with Holm-Bonferroni correction. LF-HRV (left panel) did not differ significantly between the evening and morning sessions ($F(1,81)=0.95$, $p=.33$, $\eta_p^2=0.01$) and was not influenced by sleep deprivation (vs restful sleep) in either session (main effect: $F(1,81)=0.89$, $p=.35$, $\eta_p^2=0.01$; interaction: $F(1,81)=3.54$, $p=.063$, $\eta_p^2=0.04$). HF-HRV (right panel) was also comparable between the evening and morning sessions ($F(1,81)=1.39$, $p=.24$, $\eta_p^2=0.02$) and there was no general effect of sleep deprivation (vs restful sleep; $F(1,81)=0.63$, $p=.43$, $\eta_p^2<0.01$). In the restful sleep group, HF-HRV was lower in the morning session relative to the evening session ($t=4.11$, $p<.001$; interaction: $F(1,81)=21.20$, $p<.001$, $\eta_p^2=0.21$). This effect was not observed in the sleep deprivation group ($t=2.41$, $p=.074$). N=1 participant was excluded from these analyses because their ECG data was not recorded in the morning session due to a technical fault.

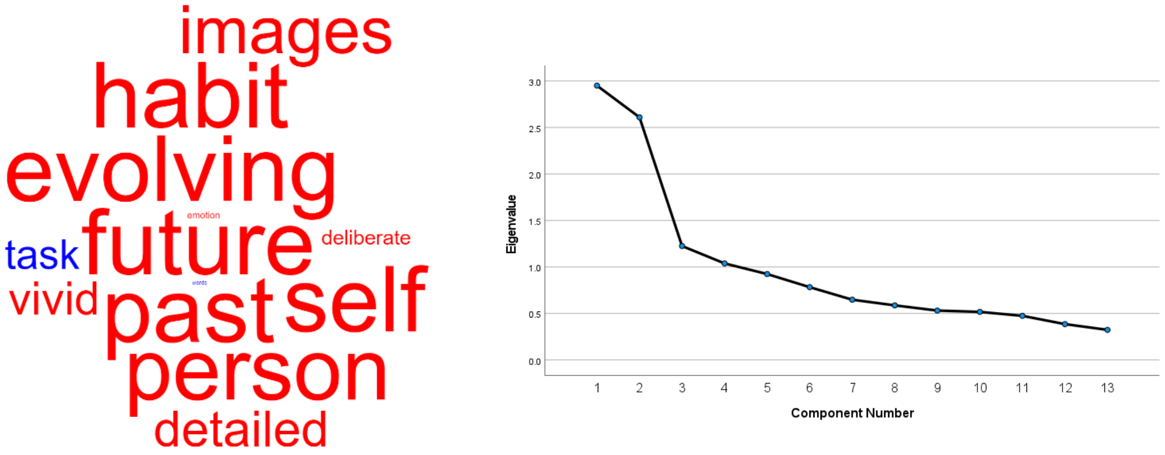


**Supplementary Figure 2.** Principal components analysis (PCA). Left: The loadings on the second component are presented as a word cloud. The colour of a word describes the direction of the relationship (red: positive, blue: negative) and the size of a word reflects the magnitude of the loading. Scores for this component were entered into a mixed-measures ANOVA with factors Task (0-back, 1-back), Session (Evening, Morning) and Group (Sleep Deprivation, Restful Sleep). There were no main effects of Task ($F$(1,80)=0.81, $p$=.37, $\eta_p^2$=0.01) or Group ($F$(1,80)<0.01, $p$=.96, $\eta_p^2$<0.01), but a main effect of Session ($F$(1,80)=3.96, $p$=.05, $\eta_p^2$=0.05) indicated that this thought pattern emerged to a greater extent in the morning than the evening. No significant interactions were observed for Task*Group ($F$(1,80)=0.06, $p$=.81, $\eta_p^2$<0.01), Session*Group ($F$(1,80)=0.50, $p$=.48, $\eta_p^2$<0.01), Task*Session ($F$(1,80)=0.27, $p$=.60, $\eta_p^2$<0.01) or Task*Session*Group ($F$(1,80)=0.15, $p$=.70, $\eta_p^2$<0.01). Right: Scree plot for the PCA.

**Supplementary Table 1.** Significant clusters found for seeds in the default mode network (DMN) and cognitive control network (CCN)

| *Seed* | *X Y Z (MNI)* | *No. voxels* | *Activation* | *Region* |
|---|---|---|---|---|
| DMN | -44 -36 58 | 1958 | Increase | Postcentral gyrus (left) |
| | 8 -18 10 | 1601 | Decrease | Thalamus (bilateral) |
| | 56 -18 46 | 1504 | Increase | Postcentral gyrus (right) |
| | 56 -30 -28 | 816 | Increase | Inferior temporal gyrus (right) |
| | -42 10 22 | 770 | Increase | Inferior frontal gyrus (left) |
| | -44 -60 2 | 728 | Increase | Lateral occipital cortex (left) |
| | -42 56 8 | 410 | Increase | Frontal pole (left) |
| | 48 16 28 | 318 | Increase | Inferior frontal gyrus (right) |
| | 6 42 -4 | 276 | Decrease | Cingulate gyrus (anterior) |
| | -68 -30 12 | 229 | Increase | Planum temporale (left) |
| | -36 -4 54 | 201 | Increase | Precental gyrus (left) |
| | 24 -2 66 | 163 | Increase | Superior frontal gyrus (right) |
| CCN | -16 42 24 | 1084 | Increase | Frontal pole (left) |

Clusters showing a significant difference in connectivity for the contrast sleep deprivation>restful sleep. Results were thresholded using Gaussian Random Field theory, with a $p<0.001$ (uncorrected, two-sided) voxel threshold and an FDR-corrected cluster threshold at $p<0.05/7=0.007$ (to also correct for testing multiple seeds)**.**

**Supplementary Table 2.** Multidimensional experience sampling (MDES) thought probes

| ***Dimension*** | ***Question*** | ***1*** | ***10*** |
|---|---|---|---|
| Task | My thoughts were focused on the task I was performing: | Not at all | Completely |
| Future | My thoughts involved future events: | Not at all | Completely |
| Past | My thoughts involved past events: | Not at all | Completely |
| Self | My thoughts involved myself: | Not at all | Completely |
| Person | My thoughts involved other people: | Not at all | Completely |
| Emotion | The emotion of my thoughts was: | Negative | Positive |
| Images | The contents of my thoughts were in the form of images: | Not at all | Completely |
| Words | The contents of my thoughts were in the form of words: | Not at all | Completely |
| Vivid | My thoughts were vivid as if I was there: | Not at all | Completely |
| Detailed | My thoughts were detailed and specific: | Not at all | Completely |
| Deliberate | My thoughts were: | Spontaneous | Deliberate |
| Habit | My thoughts had recurrent themes similar to those I have had before: | Not at all | Completely |
| Evolving | My thoughts tended to evolve in a series of steps: | Not at all | Completely |

**Supplementary Table 3.** Proportionalized affect suppression scores for negative scenes, separately for each group and memory process

| | ***Memory process*** | | |
|---|---|---|---|
| | *Baseline* | *Retrieve* | *Suppress* |
| ***Sleep-deprived*** | 0.14 (0.04) | 0.11 (0.03) | 0.13 (0.05) |
| ***Sleep-rested*** | 0.07 (0.04) | 0.17 (0.06) | 0.16 (0.07) |

Affect ratings gathered during the affect evaluation tasks were used to measure overnight changes in subjective emotional reactivity to negative scenes. Mean affect ratings were calculated for each participant, session (evening, morning), and image condition (baseline, retrieve, suppress). Affect suppression scores were calculated by subtracting the averaged values in the evening session from those in the morning session. To account for individual differences in emotional responding in the evening session, affect suppression scores were divided by the mean affect rating at the evening session to produce proportionalized affect suppression scores. Greater scores reflect more positive affect evaluations in the morning session compared with the evening session. Proportionalized affect suppression scores were applied to a mixed 2 (Group: Sleep Deprivation/Restful Sleep) x 3 (Image Condition: Retrieve/Suppress/Baseline) ANOVA. The analysis revealed no significant main effects (Group: $F(1,72)=0.01$, $p=.92$, $\eta_p^2<0.01$; Image Condition: $F(1.70,122.39)=0.88$, $p=.40$, $\eta_p^2=0.01$, *Greenhouse-Geisser corrected*), and the interaction was not significant ($F(1.70,122.39)=2.66$, $p=.082$, $\eta_p^2=0.04$, *Greehouse-Geisser corrected*).
